# A Bacteriophage Cocktail Targeting *Yersinia pestis* Provides Strong Post-Exposure Protection in a Rat Pneumonic Plague Model

**DOI:** 10.1101/2024.01.17.576055

**Authors:** Paul B. Kilgore, Jian Sha, Emily K. Hendrix, Blake H. Neil, William S. Lawrence, Jennifer E. Peel, Lauren Hittle, Joelle Woolston, Alexander Sulakvelidze, Jennifer A. Schwartz, Ashok K. Chopra

## Abstract

*Yersinia pestis*, one of the deadliest bacterial pathogens ever known, is responsible for three plague pandemics and several epidemics, with over 200 million deaths during recorded history. Due to high genomic plasticity, *Y. pestis* is amenable to genetic mutations as well as genetic engineering that can lead to the emergence or intentional development of pan-drug resistant strains. The dissemination of such *Y. pestis* strains could be catastrophic, with public health consequences far more daunting than those caused by the recent COVID-19 pandemic. Thus, there is an urgent need to develop novel, safe, and effective treatment approaches for managing *Y. pestis* infections. This includes infections by antigenically distinct strains for which vaccines, none FDA approved yet, may not be effective, and those that cannot be controlled by approved antibiotics. Lytic bacteriophages provide one such alternative approach. In this study, we examined post-exposure efficacy of a bacteriophage cocktail, YPP-401, to combat pneumonic plague caused by *Y. pestis* CO92. YPP-401 is a four-phage preparation with a 100% lytic activity against a panel of 68 genetically diverse *Y. pestis* strains. Using a pneumonic plague aerosol challenge model in gender-balanced Brown Norway rats, YPP-401 demonstrated ∼88% protection when delivered 18 hours post-exposure for each of two administration routes (i.e., intraperitoneal and intranasal) in a dose-dependent manner. Our studies suggest that YPP-401 could provide an innovative, safe, and effective approach for managing *Y. pestis* infections, including those caused by naturally occurring or intentionally developed strains that cannot be managed by vaccines in development and antibiotics.

## INTRODUCTION

*Yersinia pestis* is a Gram-negative, non-motile, coccobacillus bacterium, and human infections can lead to three deadly forms of plague, which is one of the oldest diseases responsible for the death of over 200 million people during recorded history (1, 2). Likely because of deadly combination of high fatality and contagiousness associated with pneumonic plague, *Y. pestis* was the first bacterium used as a biological warfare agent, when the Tatars catapulted plague-infected corpses into the sieged Black Sea port of Kaffa (currently Feodossia, Ukraine) in 1346 (3, 4). During World War II, the Japanese released *Y. pestis*-infected fleas to infect humans in several Chinese cities, thus causing small epidemics of plague (5). Consequently, *Y. pestis* was the key bacterium for many bioweapon development programs in the US, the former Soviet Union (FSU) (6), and other countries. Accordingly, the CDC and NIAID classified a handful of bacteria (including. *Y pestis*) and viruses as category “A” select agents (7, 8)). *Y. pestis,* now belongs to a short list of Tier-1 select agents because of its very high propensity to impact public health with mass casualties as a result of person-to-person transmission during pneumonic plague. While the incidence of plague has declined during the last several decades, cases of *Y. pestis* infections are regularly registered globally each year (9, 10). In the US, in 2015, 15 human cases of plague were reported, resulting in 4 deaths; in 2017, the island of Madagascar experienced a large outbreak, where approximately 2,350 cases of plague (∼70% in pneumonic form) occurred, including 202 deaths (2).

Current prevention and treatment modalities for *Y. pestis* infections are suboptimal at best. The current plague vaccines in preclinical development or the earlier versions of vaccines, with none approved yet by the Food and Drug Administration (FDA), have variable efficacy, limited antigenic complexity, and/or have severe side effects, also requiring regular boosting (2, 11, 12). A limited number of clinical trials are based on subunit vaccines containing only the capsular antigen F1 and the low calcium response V antigen (LcrV), with correlates of protection still undefined (13, 14). Because of the absence of a safe and effective *Y. pestis* vaccine, antibiotics remain the last bastion of defense against a possible *Y. pestis* attack or during an epidemic. However, because of bacterial high genomic plasticity (15), *Y. pestis* is amenable to genetic engineering that can lead to the emergence of pan-drug resistant strains, which would be very difficult, if not impossible, to manage using currently available antibiotics. One example of such engineering is the highly virulent, antibiotic-resistant F1^-^ strain(s) developed in the Institute of Applied Microbiology (located in Obolensk, Russia) (16). Other antibiotic-resistant mutants may emerge or be similarly engineered by other countries and/or terrorist organizations, which could be intentionally or unintentionally released. The consequences associated with the dissemination of *Y. pestis* strains that are untreatable with antibiotics could be catastrophic. Thus, there is a critical need to develop novel, safe, and effective treatment approaches for managing *Y. pestis*-associated diseases, including those caused by antigenically distinct strains (against which vaccines may not be optimally effective) and by those that cannot be killed by available antibiotics. Lytic bacteriophages can provide one such alternative approach.

Bacteriophages (or phages) are viruses that target bacterial hosts. They were identified during the early part of the 20^th^ century by Felix d’Herelle (17, 18) and were first used successfully to treat bubonic plague in 1925 (19). Phages are the oldest (∼3-billion-years-old) and most ubiquitous organisms (∼10^30^-10^31^ phage particles) on Earth. Lytic phages are very effective at killing their targeted host bacteria and, in contrast to antibiotics, they are very specific (i.e., phages will lyse related strains or a subgroup of strains, but will not lyse strains of other, unrelated bacteria). During the early 20^th^ century, their remarkable antibacterial activity prompted the use of phages for treating many bacterial diseases, but for various reasons (reviewed in (20)), their use gradually declined in the West after antibiotics became widely available. However, the emergence of multi-antibiotic-resistant pathogens has rekindled interest in the possible therapeutic applications of bacteriophages. In this context, the mechanisms by which antibiotics and lytic phages kill bacteria (and the mechanisms of bacterial resistance to antibiotics vs. phages) are fundamentally different (21). As a result, lytic phages can kill bacteria that cannot be eliminated by currently available antibiotics. Further, the development of an antibiotic resistance does not impact the lytic potency of phages and vice-versa. Thus, lytic phages may provide a critical complementing modality to antibiotics, for preventing and/or treating diseases caused by multi-antibiotic-resistant bacteria.

Several previous studies have shown *in vitro* phage efficacy against *Y. pestis* (22–24); however, a few studies have demonstrated *in vivo* efficacy (25, 26). In one study, phage treatment was shown to delay progression of pneumonic plague; however, all mice still succumbed unless antibiotic therapy was co-administered (26). Here, we assessed a four-phage preparation, YPP-401, to target *Y. pestis in vitro* and *in vivo*. YPP-401 demonstrated 100% *in vitro* lytic activity against all *Y. pestis* strains tested, including the virulent, wild-type *Y. pestis* CO92. YPP-401 also provided post-exposure protection *in vivo* in our established Brown Norway rat aerosol challenge model (27) using different treatment strategies. Phage treatment was initiated at either “early” (i.e., 18h post-infection [hpi]) or “late” (i.e., 42hpi) times post-exposure with different dosing regimens. When phage treatment was initiated at 18hpi, both intranasal (i.n.) and intraperitoneal (i.p.) administrations were effective at treating pneumonic plague in a dose dependent manner. This study demonstrates YPP-401 as a promising treatment modality to combat pneumonic plague, the deadliest form of the disease.

## RESULTS

### The bacteriophage cocktail YPP-401 and its component monophages demonstrate host-specific lytic activity *in vitro*

A proprietary bacteriophage cocktail formulation, YPP-401, containing 4 monophages (i.e., YPIX-2, -4, -5 and -6) was developed and characterized using a combination of previously identified phages specific for *Y. pestis* (23, 28). The YPP-401 component monophages previously were also shown to have 100% lytic activity against all tested high containment/select agent *Y. pestis* strains (23, 28).

Here, we assessed the ability of the YPP-401 formulation to target BSL-2 *Yersinia* spp. (**Table 1**) and non-*Yersinia* cultures (**Figure S1**) *in vitro* using a combination of liquid and/or spot test (29) assays at various phage concentrations and bacterial growth conditions. Additionally, the virulent wild-type *Y. pestis* CO92 strain (i.e., the challenge strain) was susceptible to the YPP-401 cocktail along with all tested *Y. pestis* and *Y. pseudotuberculosis* strains. Some *Y. pestis* phages have been reported to have activity in laboratory medium but not blood (22); therefore, we also performed the plate-based assays for *Y. pestis* CO92 susceptibility on blood agar plates and obtained similar results (data not shown). Only one *Y. aldovae* strain was susceptible to the YPP-401 formulation, while strains representing 4 other *Yersinia* spp. (i.e., *Y. enterocolitica*, *Y. frederiksenii*, *Y*. *kristensenii*, and *Y*. *mollaretii*) were not susceptible. Additionally, to demonstrate the specificity of YPP-401 for *Yersinia* spp., susceptibility to the YPP-401 cocktail was evaluated against a representative panel of bacteria (**Figure S1**). All of the non-*Yersinia* spp. tested were not susceptible to YPP-401 with the exception of several *E. coli* strains. As expected, YPP-401 was highly specific for *Yersinia* spp. compared to antibiotic treatments, including ciprofloxacin, a first-line antibiotic approved for treatment of plague that broadly inhibited growth of various bacteria (**Figure S1**).

**Table 1.** In Vitro Lytic Activity of YPP-401 Against Yersinia spp.

| Genus | Species | Strain | Susceptible* |
| --- | --- | --- | --- |
| *Yersinia* | *aldovae* | ATCC 35236 | + |
| *Yersinia* | *aldovae* | CNY 7112 | - |
| *Yersinia* | *enterocolitica* | 30.42.67 | - |
| *Yersinia* | *enterocolitica* | 8081 | - |
| *Yersinia* | *frederiksenii* | CNY 867 | - |
| *Yersinia* | *frederiksenii* | WA 935 | - |
| *Yersinia* | *kristensenii* | ATCC 33638 | - |
| *Yersinia* | *kristensenii* | WE 180/98 | - |
| *Yersinia* | *mollaretii* | WE 149/92 | - |
| *Yersinia* | *mollaretii* | WS 43/94 | - |
| *Yersinia* | *pestis* | A12 D6 | + |
| *Yersinia* | *pestis* | A1122 | + |
| ***Yersinia*** | ***pestis*** | **CO92** | + |
| *Yersinia* | *pestis* | Harbin 35 | + |
| *Yersinia* | *pestis* | K25 D80 | + |
| *Yersinia* | *pestis* | KIM D2 | + |
| *Yersinia* | *pestis* | KIM D19 | + |
| *Yersinia* | *pestis* | KIM D23 | + |
| *Yersinia* | *pestis* | KIM D27 | + |
| *Yersinia* | *pestis* | Kimberly D13 | + |
| *Yersinia* | *pestis* | KUMA D8 | + |
| *Yersinia* | *pestis* | TS D4 | + |
| *Yersinia* | *pestis* | Yokohama D11 | + |
| *Yersinia* | *pseudotuberculosis* | ATCC 23207 | + |
| *Yersinia* | *pseudotuberculosis* | ATCC 29833 | + |
| *Yersinia* | *pseudotuberculosis* | CDC 542-84 | + |
| *Yersinia* | *pseudotuberculosis* | CDC 801-84 | + |

\*Susceptibility tested across various phage concentrations in liquid and/or solid spot test assays; **BOLD**, virulent, wild-type *Y. pestis* strain used in challenge studies.

### YPP-401 provides protective efficacy *in vivo* when treatment is initiated at 18h post-exposure

Previous studies conducted by our laboratory using the aerosol challenge model of pneumonic plague in Brown Norway rats implanted with telemetry showed that the febrile response (≥39°C temperature for at least 1h of duration) usually started after 36 hpi (unpublished data). Here, animals in cohort 1 were subjected to a full body aerosol challenge (*Y. pestis* CO92 with a presented dose (Dp) of 9.56 x 10^6^ to 1.24 x 10^7^ CFU) followed by a YPP-401 treatment regimen starting at 18 hpi as depicted in **Figure 1A**. At 18 hpi, cohort 1 animals were administered 4 total doses of YPP-401 over 24-42 h approximately every 6-12 h (i.e., 18, 24, 30 and 42 hpi) by one of 3 separate routes: oral (∼2×10^10^ plaque-forming units [PFU]/dose); i.n. (∼4×10^9^ PFU/dose); or i.p. (∼2×10^10^ PFU/dose) (**Figure 1A**). Control treatment groups included vehicle alone (i.e., PBS) delivered i.p. 4 times at the same intervals as YPP-401. Levofloxacin (50 mg/kg) was delivered orally, daily for 10 days. Survival data for cohort 1 is shown in **Figure 1C**. Rats treated orally with YPP-401 succumbed to infection within 2-6 days, with 2 animals demonstrating a delayed time to death compared to controls. Intraperitoneal treatment with YPP-401 resulted in 87.5% survival with one rat succumbing to disease at day 5 post-infection (dpi). Intranasal treatment with YPP-401 resulted in 50% survival, with rats succumbing to infection on 4-9 dpi. Due to administration volume limitations for i.n. delivery, animals received 1/5^th^ of the YPP-401 dose than that delivered by the i.p. route. All vehicle treated mice succumbed to infection by 3 dpi. Rats treated with levofloxacin, the positive control, had 100% survival at 14 dpi after 10 daily doses of the antibiotic.

**Figure 1.**
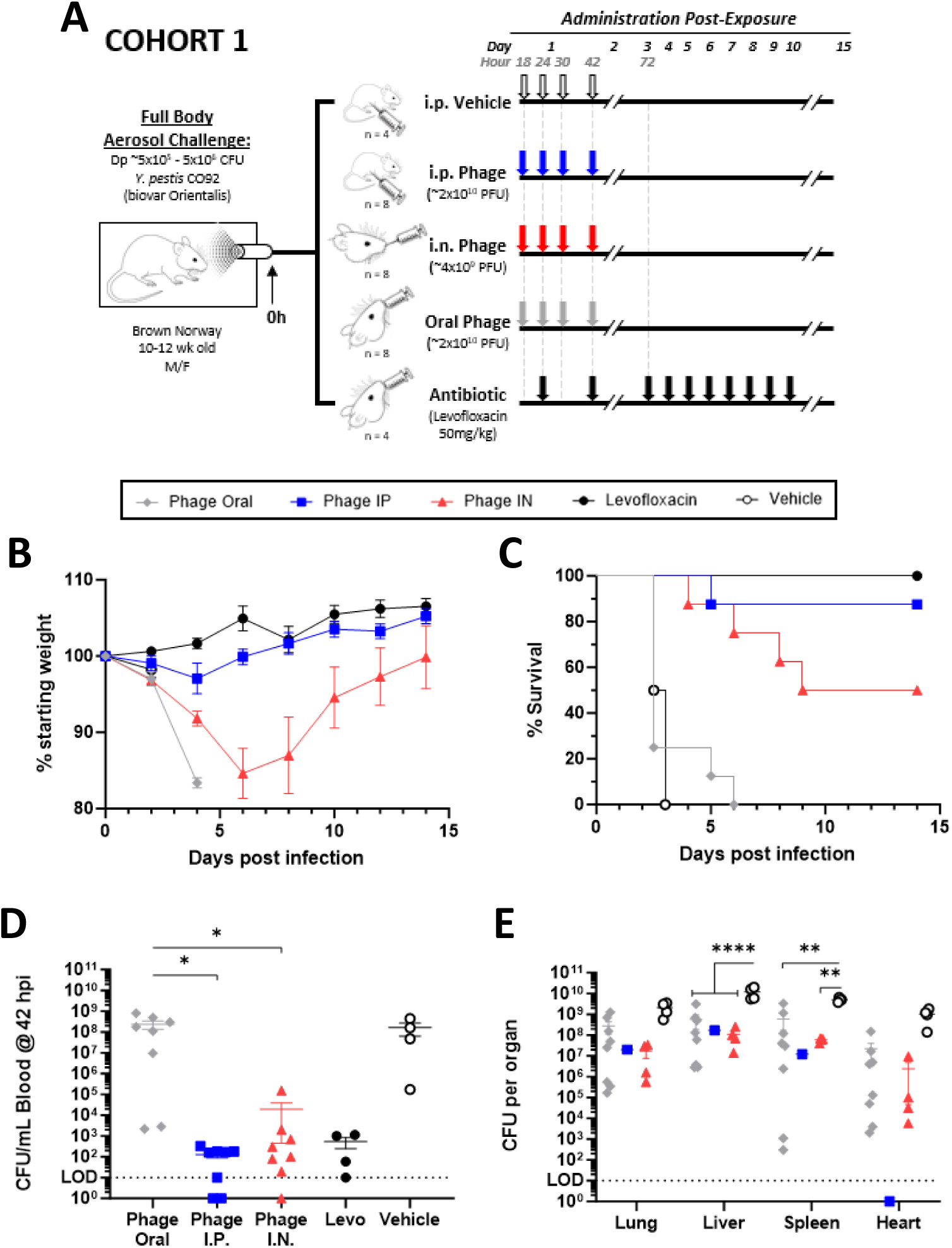
YPP-401 provides post-exposure efficacy against *Y. pestis* in rats when delivered intraperitoneally or intranasally at 18 hpi. <u>COHORT 1</u>: **(A)** Schematic of study design and dosing regimens. Shown are **(B)** body weight,**(C)** survival and (D) *Y. pestis* burden in the blood at 42 hpi (taken immediately prior to 42h treatment timepoint). Other measures were **(E)** terminal organ bacterial burden (i.e., lung, liver, heart, spleen), and clinical score (i.e., appearance, activity, respirations, facial expression; **Table S2**). Brown Norway rats (M/F; 10-12 wk) received a full body aerosol challenge (0 h) of WT virulent *Y. pestis* strain CO92 at Dp ∼9.56 x 10^6^ to 1.24 x 10^7^ CFU. At 18 hpi, cohort 1 received either 2 days (total 4 doses, BID, ∼q6-12h) of i.p. vehicle (white/open circles; PBS; n=4), i.p. YPP-401 (blue squares; ∼2×10^10^ PFUs; n=8), i.n. YPP-401 (red triangles; ∼4×10^9^ PFUs; n=8), oral YPP-401 (gray diamonds; ∼2×10^10^ PFUs; n=8), or 10 oral QD doses of levofloxacin (levo) (black circles; 50 mg/kg; n=4). *p<0.05; **p<0.01; ***p<0.001. Studies were performed at UTMB under ABSL-3.

### Clinical scores and bacterial burden in blood and organs correlate with survival in YPP-401 treated animals

Clinical signs and symptoms were assessed and included weight loss (**Figure 1B**) and clinical scores (**Table S2**) that accounted for independent variables for appearance, activity level, respiration, and facial expression. In general, across all groups, clinical scores correlated with time to death (**Table S2**). Additionally, in rats receiving i.n. YPP-401, 7 out of 8 rats lost 15-25% bodyweight with peak weight loss occurring by 6-8dpi. Surviving animals began recovering weight by 6-10 dpi (**Figure 1B**). In contrast, a significant weight loss in the i.p. YPP-401 treated rats was only observed in the single animal that succumbed to disease at 5 dpi (**Figure 1B**). These results are consistent with our previous studies (unpublished data) where we typically did not observe significant weight loss until 4 dpi, and thus explains the minimally observed weight loss in the vehicle- and YPP-401 orally-treated animals that succumbed to disease within 4 dpi (**Figure 1A**).

To determine the effect of YPP-401 treatment on bacteremia, blood was collected at 42 hpi (i.e., immediately before the last phage dose) and titrated on SBA plates. Groups with low survival rates (i.e., vehicle and oral YPP-401) had high levels of *Y. pestis* in the blood with an average of 10^8^ CFU/mL (**Figure 1D**). Groups treated with 4 doses of YPP-401 by i.p. or i.n. had significantly lower levels of *Y. pestis* in the blood than animals treated orally (**Figure 1D**). Intraperitoneal YPP-401 treated rats had similar levels of *Y. pestis* bacteremia as the levofloxacin treated controls (**Figure 1D**).

Total organ bacterial burden also was evaluated in post-treatment animals that succumbed to disease. Bacterial loads in the terminal lungs, liver, spleen, and heart were quantitated from tissue homogenates on SBA plates (**Figure 1E**, **Table S2**). As expected, the vehicle treated rats had the highest bacterial load in all 4 organs analyzed. Rats treated with YPP-401 that succumbed, regardless of the administration route, had lower levels of *Y. pestis* in the lungs, liver and spleen compared to the vehicle controls, and generally had similar levels among all YPP-401 treated groups. Two animals treated orally with YPP-401 had significantly lower levels of *Y. pestis* in the blood and correlated with delayed time to death (**Table S2**). In the one i.p. YPP-401 treated animal that succumbed to disease, only 180 CFU/ml of blood were detectable and there was a delayed time to death (**Table S2**). The oral and i.n. YPP-401 treated groups had similar levels of *Y. pestis* in the heart. In general, compared to animals that succumbed to disease, YPP-401 treated rats had statistically lower levels of *Y. pestis* in blood and organs compared to the vehicle control (**Figure 1D** and **1E**, **Table S2**).

### YPP-401 provides dose-dependent protective efficacy and reduced bacterial loads *in vivo* when delivered intranasally beginning at 18h post-exposure

To determine whether increasing the i.n. YPP-401 dose could improve post-exposure protection when delivered starting at 18hpi, cohort 2 animals were challenged with *Y. pestis* CO92 by the aerosol route as described above for cohort 1 (**Figure 1A**). Here, rats treated with 4 total doses of YPP-401 (between 18 and 42 hpi) were given approximately 5X the dose (i.e., ∼2×10^10^ PFU/dose) delivered in cohort 1 (i.e., (∼4×10^9^ PFU/dose). As in cohort 1, rats were randomly distributed into 4 aerosol challenge runs where the presented dose between runs ranged from 9.79 x 10^6^ to 1.06 x 10^7^ CFUs.

**Figure 2B** shows that increasing the YPP-401 i.n. dose increased survival from 50% in cohort 1 (**Figure 1C**) to 87.5% (**Figure 2B**). Control groups behaved as expected with all vehicle treated animals succumbing by 3 dpi and all levofloxacin treated rats, which received 10 daily doses, surviving through 14 dpi (**Figure 2B**). These results, interestingly, demonstrate that i.n. YPP-401 treatment can be as effective as i.p. YPP-401 treatment when administered at the same dose. Cohort 2 rats treated with YPP-401 i.n. lost no more than ∼10% body weight (**Figure 2A**) compared to cohort 1 YPP-401 i.n. treated animals (**Figure 1B**), with peak weight loss observed at 6 dpi, and all surviving rats gaining weight by day 8 (**Figure 2A**). Levofloxacin treated rats lost minimal weight with one rat losing 5% body weight by 2 dpi, and which recovered to pre-infection levels by 4 dpi (**Figure 2A**).

**Figure 2.**
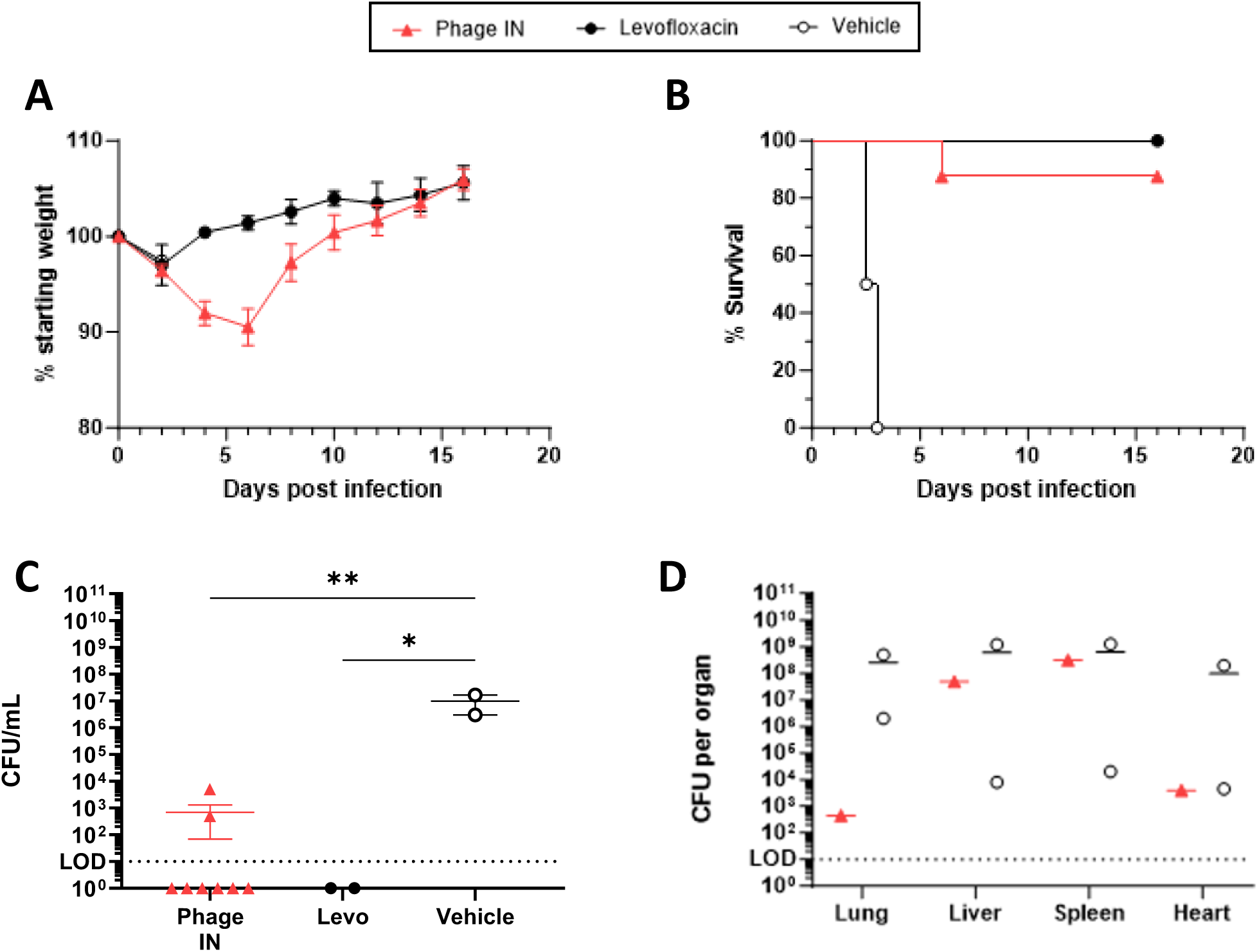
YPP-401 provides dose-dependent post-exposure efficacy against *Y. pestis* in rats when delivered intranasally at 18 hpi. <u>COHORT 2</u>: Animals were challenged as described in Figure 1A. Similar dosing regimens starting at 18 hpi were used with the following changes: cohort 2 received either 2 days (total 4 doses, BID, ∼q6-12h) of i.p. vehicle (white/open circles; PBS; n=2), i.n. YPP-401 (red triangles; ∼2×10^10^ PFUs; n=8), or 10 oral QD doses of levo (black circles; 50 mg/kg; n=2). Shown are **(A)** body weights, **(B)** survival, and (C) *Y. pestis* burden in the blood at 42 hpi. (taken immediately prior to 42h treatment timepoint). Other measures were **(D)** terminal organ bacterial burden (i.e., lung, liver, heart, spleen), and clinical score (i.e., appearance, activity, respirations, facial expression; **Table S3**). *p<0.05; **p<0.01; ***p<0.001. Studies were performed at UTMB under ABSL-3.

As performed for cohort 1 above, bacteremia (**Figure 2C**, **Table S3**) and organ burden (**Figure 2D**, **Table S3**) were assessed at either 42 hpi or terminally, respectively. Seventy-five percent (6/8) of YPP-401 i.n. treated rats had no detectable *Y. pestis* in the blood (**Figure 2C**, **Table S3**). The 2 animals that had detectable bacteria in the blood were at a significantly reduced level (i.e., <10^4^ CFU/mL) compared to the vehicle controls (i.e., >10^6^ CFU/mL; **Figure 2C**) and were similar to levels in cohort 1 animals treated with levofloxacin or YPP-401 delivered by i.n. or i.p. routes (**Figure 1D**). The cohort 2 levofloxacin treated rats (n = 2) had no detectable *Y. pestis* in the blood. Total organ burden in cohort 2 rats that succumbed was evaluated; however, due to the high survival rates, statistical methods could not be applied. *Y. pestis* found in the lungs of the only terminal YPP-401 i.n. treated rat was 4-6 logs lower than vehicle treated rats (**Figure 2D**), while no statistically significant conclusion could be reached for bacterial burden in the liver, spleen, and heart compared to the vehicle controls due to the low number of animals (n = 2) and high variability in the vehicle control group (**Figure 2D**).

### Highly protective YPP-401 dosing regimens initiated at 18h post-exposure are not sufficient to protect against *Y. pestis* challenge at a more delayed time post-exposure when clinical signs and symptoms are apparent

To determine whether the YPP-401 i.p. protective post-exposure dosing regimen used in cohort 1 (**Figure 1A**) is sufficient to protect animals at a more delayed time post-exposure, a third cohort (cohort 3) was treated starting at 42 hpi after *Y. pestis* CO92 aerosol challenge. It should be noted, as has been shown previously, that at 42 hpi, clinical signs and symptoms of disease are typically apparent and even frontline antibiotic therapy initiated at this timepoint has much reduced efficacy (30, 31). As shown in **Figure 3B**, rats treated with YPP-401 i.p. starting at 42 hpi, planned to be administered 6-12 h apart, were only able to receive 1 or 2 doses (**Table S4**) of YPP-401 before succumbing to disease, and thus was ineffective at providing protection under these conditions. Moreover, even the levofloxacin control treatment, which reliably rescues 100% of treated rats when administered at 18 hpi, was only effective in 50% of the animals (n = 2) with one rat succumbing to infection having received 1 out of the 10 intended daily doses (**Figure 3B**, **Table S4**). Under emergency or mass treatment settings, i.p. administration is not practical; therefore, we also assessed whether YPP-401 delivered intramuscularly (i.m.) could provide post-exposure protection under these delayed and stringent conditions where treatment was initiated at 42 hpi. As seen with YPP-401 delivered i.p., i.m. administration did not improve protection under this dosing regimen (**Figure 3B**, **Table S4**).

**Figure 3.**
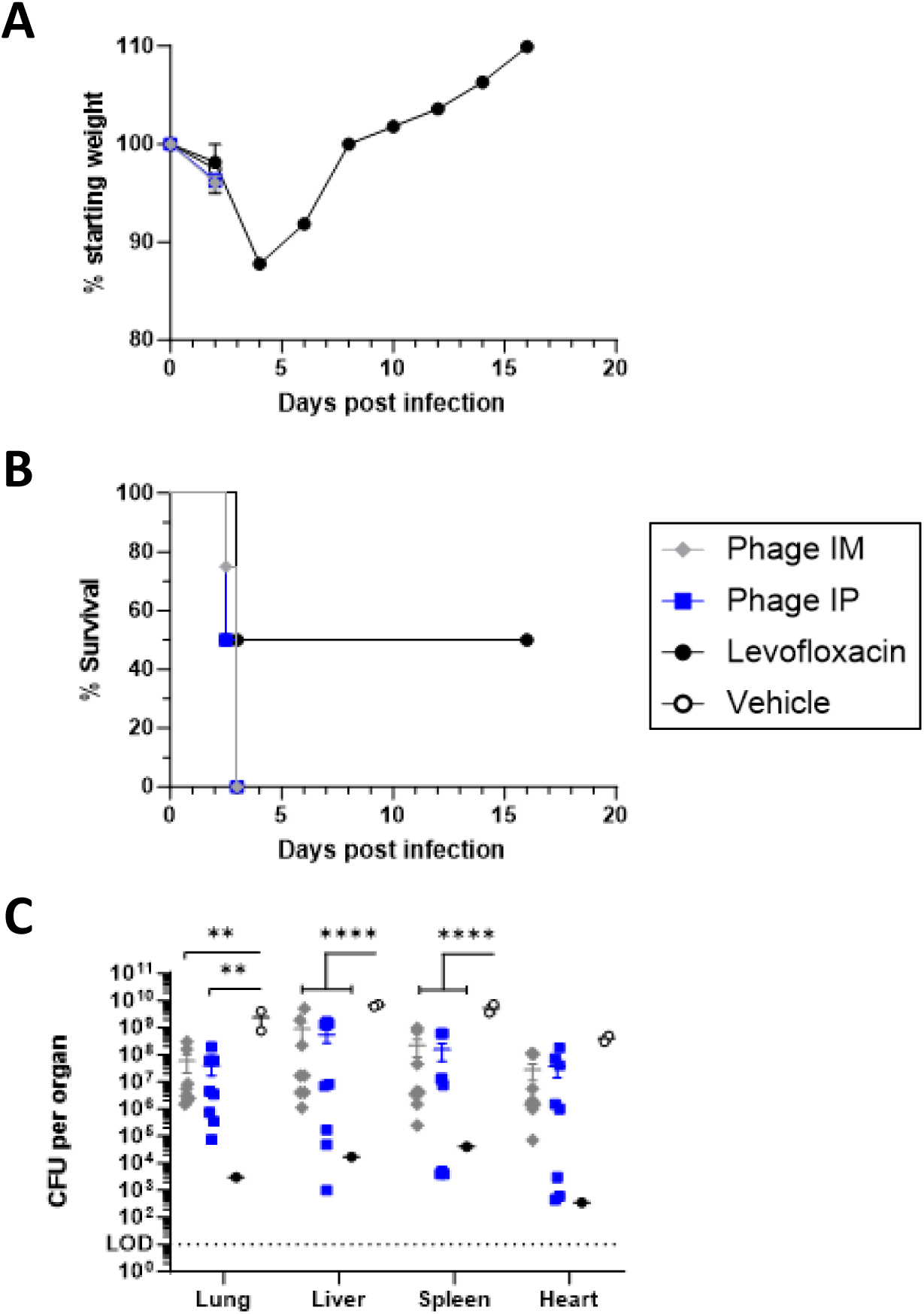
YPP-401 dosing regimens protective at 18 hpi are insufficient when initiated at 42 hpi. <u>COHORT 3</u>: Animals were challenged as described in Figure 1A. Similar dosing regimens were started at 42 hpi with the following changes: cohort 3 received either 1 or 2 doses (BID, ∼q6-12h) of i.p. vehicle (white/open circles; PBS; n=2), i.m. YPP-401 (gray circles; ∼5×10^10^ PFUs; n=8), i.p. YPP-401 (blue squares; ∼5×10^10^ PFUs; n=8), or 1 or 10 QD doses of levo (black circles; 50 mg/kg; n=2). Shown are **(A)** body weights, **(B)** survival, and **(C)** *Y. pestis* burden in terminal organs (i.e., lung, liver, heart, spleen), and clinical score (i.e., appearance, activity, respirations, facial expression; **Table S4**). *p<0.05; **p<0.01; ***p<0.001. Studies were performed at UTMB under ABSL-3.

Prior to treatment at 42 hpi, blood was collected to evaluate levels of bacteremia. All rats had high levels (up to 10^7^ CFU/mL) of *Y. pestis* in the blood (**Table S4**). Rats that succumbed were necropsied to assess the levels of *Y. pestis* present in the lungs, liver, spleen, and heart. While vehicle treated rats had uniformly high levels of *Y. pestis* in these organs at levels >10^8^ CFU/organ, the number of *Y. pestis* found in YPP-401 or levofloxacin treated rats were more varied (**Figure 3C**, **Table S4**) likely due to killing either by phage or antibiotics, respectively. The single levofloxacin treated rat that survived showed enhanced morbidity over that seen in cohorts 1 and 2 with >12.5% peak body weight loss by 4 dpi with recovery occurring by 6 dpi (**Figure 3A**). **Table S4** shows the values for individual rats that succumbed to infection (e.g., the number of treatment doses received, bacteremia, organ burden, clinical score before succumbing to disease, and time to death).

## DISCUSSION

### *Yersinia pestis*, a potential biotreat

The diseases *Yersinia pestis* cause have a high fatality rate of approximately 30% (bubonic plague), and with septicemic and pneumonic plague, if not treated within 24 h of symptoms onset, has a mortality rate approaching 100% (32). The bacteria can spread rapidly person-to-person via the aerosol route, which makes containment difficult. According to some early estimates from the World Health Organization (33), 50 kg of *Y. pestis* released as an aerosol over a city of about 5 million people will cause pneumonic plague in about 150,000 inhabitants. Of these infected people, approximately 36,000 would be expected to die within a few days. The report also suggested that a significant number of the city’s inhabitants would attempt to flee, further spreading the disease and contributing to the escalating chaos. The validity of the latter suggestion, and the profound psychological impact that the diagnosis of plague can have on a community, was confirmed in Surat, India in 1994, when >600,000 of the estimated population of 2 million people fled the city after early reports of pneumonic plague cases (34). The strain of *Y. pestis* causing the Surat outbreak was antibiotic sensitive and could be effectively controlled with proper antibiotic therapy.

The consequences of similar outbreaks will be even more destructive if the outbreak-causing strain is multidrug resistant (e.g., naturally mutated or genetically engineered to be resistant to all currently available antibiotics by a terrorist organization or rogue nation). Thus, we may lack effective tools for preventing and/or treating the diseases the organism causes. In this context, although multidrug resistance has been relatively rare in *Y. pestis*, isolates of *Y. pestis* showing tetracycline resistance and reduced susceptibility to streptomycin have been described in some African countries and China. Likewise, multidrug-resistant isolates have been reported from Madagascar, and a strain resistant to most of the antibiotics commonly used in prophylaxis and therapy of plague was isolated in Mongolia (35–38). Moreover, several countries, including the former Soviet Union, used to have bioweapons programs designed to engineer *Y. pestis* clones that would be more difficult to treat with antibiotics and could also be better aerosolized for even larger and more rapid spread (6).

While those programs supposedly have been abandoned, there is a real possibility that some of them continue(d) to operate secretly and/or previously developed multidrug resistant, super aerosolizing *Y. pestis* strains could be obtained and intentionally released by bioterrorists. The consequences of such an attack would be devastating. Of note, in November 2021, Microsoft founder Bill Gates highlighted the risk of bioterrorism, emphasizing that the consequences could be far worse than those caused by the COVID-19 pandemic, urging world leaders to allocate “tens of billions” for research and development to prepare for such attacks and the next pandemic (39). This would be a very serious situation because of the degree of devastation bioterrorism attacks can cause. Although preventing bioterrorism remains the optimal choice for avoiding attack-associated casualties and loss of civil services, it is unlikely that all bioterrorism attacks can be prevented. Thus, it is important to have the infrastructure and tools to deal effectively with the consequences of such attacks, including minimizing human casualties and infrastructure damage.

### Phage therapy, a promising approach to mitigate bacterial infections

There are multiple favorable qualities in support of therapeutic phages. Phages, in contrast to antibiotics and other antimicrobial agents, are extremely host specific (i.e., they attack specific bacterial strains or subgroups of strains) and thus have minimal potential for the disruption of normal or commensal flora. Lytic phages are bactericidal, as opposed to bacteriostatic, thereby possibly providing faster and unreversible killing of the pathogen. The mechanism of action for phages is independent from traditional antimicrobial mechanisms and thereby can be used against multi-drug resistant (MDR) strains (40, 41). Finally, there is a long history of lack of toxicity with phages, and they have even been used in severely immunodeficient humans (42–44).

The first ever clinical use of bacteriophages was successfully conducted in France in 1919 (45). In fact, the first therapeutic application of phages for bubonic plague was performed successfully by d’Herelle in Egypt in 1925 in four patients (46). Before antibiotics became available, phages were perhaps the most frequently and successfully used therapies for treating human bacterial infections (reviewed by (20, 47)). While in the pre-antibiotic era, phage-based therapies were used to treat infections, they often produced inconsistent results. Several factors likely contributed to the unpredictable results of these very early phage therapies (reviewed in detail (41, 48–50)) and include: poor understanding of phage biology, inadequate diagnostic bacteriology techniques, poor characterization of the phage products, and variable purities of the phage preparations. During the rise of the antibiotic era, phage therapies fell out-of-favor in the U.S. and in Western Europe, but continued to be used in the former Soviet Union and in Eastern Europe, and is standard of care for various bacterial indications even up to today. The semi-anecdotal data (e.g., due to lack of well controlled clinical trials likely resulting from their well accepted and widespread use in these countries) that exists from their ongoing use in these regions suggests that phages, when carefully selected, are often more effective than antibiotics alone in treating many infections (e.g., *Staphylococcus aureus* infections of the lung parenchyma and pleura) (51–53). Significantly, even when disease-causing bacteria were antibiotic-sensitive, administering phages together with traditional antibiotics was reported to improve the clinical outcome (reviewed in (20)). Although the underlying mechanisms for this phenomenon have not been rigorously elucidated, it is likely that simultaneously exposing a pathogen to two antibacterial agents possessing different modes of actions (i.e., phages and antibiotics) resulted in more effective bacterial killing than did treatment with just the traditional antibiotic alone (26). Indeed in this study, a combination of phage and antibiotic therapy was able to achieve complete protection when neither therapy was effective alone.

### YPP-401 is a potent phage-based product that targets *Y. pestis*

We have identified a candidate *Y. pestis* phage cocktail preparation, YPP-401. Using proprietary software PhageSelector™ (developed by Intralytix), which facilitates the design of optimally effective phage preparations, we formulated the candidate YPP-401 for *Y. pestis.* The program uses various algorithms to analyze the database of bacterial strains in conjunction with phage sensitivity data, in order to suggest which phages should be included in the preparations. For example, the program can examine the data for phage potency variation (i.e., it can determine the total number of bacterial strains that each phage is capable of lysing). This method promotes redundancy by identifying phages that have high-kill counts and, therefore, are most likely to lyse the same bacterial strains lysed by other high-kill count phages. This redundancy in killing not only increases the potency of the cocktail, but also allows for little to no phage resistance development (54–56).

The program can also perform analyses to determine the bacterial lysis efficiency, which orders the candidate phages according to their ability to lyse the maximum number of bacterial strains listed in the bacterial database (i.e., it identifies the phages that are most diverse in their lytic activity against the strains listed in the database). The program can incorporate other available parameters that would further facilitate designing optimally effective phage preparations, such as **1)** the burst size of each candidate phage where the larger the burst size, the stronger the lytic potency of a phage may be; **2)** the genomic composition of each candidate phage; and **3)** phages that attach to different receptors on the bacterial host’s surface may enhance the phage cocktail’s lytic potency and reduce the risk of bacterial resistance developing against the cocktail. Using PhageSelector™, we identified the optimal combination of *Y. pestis* phages to be included in our candidate YPP-401 preparation. This potent phage cocktail contains 4 lytic bacteriophages with excellent lytic potency against all tested *Y. pestis* strains (**Table 1** and (23, 28)), as determined using the classical Spot Test method ((57)). This collection includes geographically (including the US, Iran, Georgia, Azerbaijan, Armenia, Africa, and Japan) and temporally (over > 30 years) distinct strains collected from various sources (e.g., humans, rodents, fleas, soil).

### YPP-401 provides strong post-exposure protection in a rat challenge model

While several published studies have shown phages can target *Y. pestis in vitro*, only a few studies have examined *in vivo* efficacy of *Y. pestis* phages (22, 23, 26). To this end, we examined whether YPP-401 could provide post-exposure protection at “early” (i.e., 18 hpi; **Figure 1** and **2**) or “late” (i.e., 42 hpi; **Figure 3**) times post-lethal challenge with *Y. pestis* CO92 using different dosing regimens, varying route (i.e., i.p., i.n., p.o., i.m.), and/or YPP-401 dose (∼4×10^9^-5×10^10^ PFU/dose) in a previously characterized, rat aerosol challenge model (27, 31, 58–60).

In cohort 1 (**Figure 1A**), rats received a “low” dose of YPP-401 (∼2×10^10^ PFU/dose by i.p. or p.o., or ∼4×10^9^ PFU/dose by i.n. – dose varied due to maximum amount of volume delivered per route) starting at 18h post-challenge. YPP-401 was administered about every (q) 6-12h for a total of 4 doses until 42 hpi, and the time at which animals typically become symptomatic. Control groups included vehicle alone or levofloxacin (levo; 50 mg/kg), a first line antibiotic approved for the treatment of pneumonic plague, administered p.o. daily (i.e., QD) for 10 days and previously known to fully protect in this model. Rats receiving low dose YPP-401 by i.p. had an 88% survival rate while those treated by i.n. had a 50% rate of survival (**Figure 1C**). Despite a slight delay in mortality, those receiving YPP-401 p.o. succumbed. Rats receiving YPP-401 i.p. had similar levels of *Y. pestis* in their blood as the levofloxacin controls at 42 hpi (**Figure 1D**). Bioavailability of phage by the oral route is likely poor. In an earlier study (61), it was reported that while orally administered phages could be detected in the blood and liver, there was a 4-log less number of phages compared to the numbers administered (61). Bioavailability has also been compared between i.p. and i.n. phage therapy. While i.p. adminsteration of phages led to their higher numbers in the spleens of treated animals, i.n. therapy resulted in more phages in the lungs by 3-4 logs (26).

To determine whether i.n. YPP-401 post-exposure protection could be improved by increasing the YPP-401 dose, rats in cohort 2 received an intermediate dose (∼2×10^10^ PFU/dose, ∼5X the cohort 1 i.n. dose) of YPP-401 starting at 18h post-challenge and ∼q6-12h for a total of 4 doses until 42 hpi (**Figure 2**). We showed a dose dependent effect on survival after only 4 intermediate post-exposure doses of YPP-401 where the rate of survival increased to 88% (**Figure 2B**) compared to 50% in the low dose i.n. group in cohort 1 (**Figure 1C**). In another study using 1×10^9^ PFU of a different phage cocktail, no efficacy with phage therapy was observed, suggesting a dose-dependent effectiveness for treatment of pneumonic plague (26). Stringency of test parameters was increased further by assessing post-exposure efficacy at “late” times post-challenge (i.e., 42 hpi, after animals become symptomatic and a time where existing antibiotics have suboptimal efficacy; **Figure 3**). “Late” post-exposure treatment with levofloxacin dosed daily for 10 days starting at 42 hpi, provided a suboptimal survival rate of 50%. When YPP-401 was delivered starting at 42 hpi by i.p. or i.m. at 2X the cohort 1 dose, all animals succumbed; however, only 1 or 2 phage doses were able to be delivered prior to time of death (**Table S4**).

It has been well documented that the timing of treatment for pneumonic plague is very important. Most literature suggests that if not treated within 24 h of symptom onset, pneumonic plague is almost uniformly fatal (9, 62, 63). As shown in this study, levofloxacin treatment initiated at 18 hpi is very effective at completely protecting rats from pneumonic plague. Byrne, et al. also have shown that Ciprofloxacin, Streptomycin, or several other antibiotic treatments are 100% effective when administered 24 hpi in mice that were challenged with 100 LD_50_ of *Y. pestis* via the aerosol route. When treatment was delayed until 42-48 hpi, the efficacy dropped and was varied (30). At lower challenge doses of 12 LD_50_ using an i.n. infection rat model instead of aerosol infection, and delaying levofloxacin treatment until 42 hpi had no deleterious impact on survival (31). However, when Cethromycin was used as the treatment, at 30 LD_50_ in the i.n. rat challenge model, delaying treatment past 24 hpi reduced survival from 100% at 24 hpi to 40% when treatment was started at 36h or 48h post-infection (31). The same study by Byrne, et al. showed that when treatment was delayed in mice from 24 to 42 hpi, protection rates dropped to 62% and 59% for Ciprofloxacin and Streptomycin, respectively (30). In the studies we presented here, rats were challenged with ∼7,000 LD_50_ via the aerosol route; thus, it is not surprising that levofloxacin treatment only protected 50% of the rats and that the 1-2 doses of YPP-401 delivered starting at 42 hpi were suboptimal.

Overall, these data demonstrate the promise of YPP-401 as a post-exposure treatment option to manage *Y. pestis* pneumonic infections. These studies are the first reported standalone use of effective phage therapy in an aerosol challenge model of pneumonic plague using rats. It provides promising data for future studies that could help usher phage therapy into the clinic for treatment of pneumonic plague.

## MATERIALS AND METHODS

### Bacterial Strains

Twelve avirulent *Y. pestis* isolates (**Table 1**) were obtained through BEI Resources, Manassas, VA (supported by the National Institute of Health [NIH]/National Institute of Allergy and Infectious Diseases [NIAID]). These 12 isolates are each lacking at least one of the four features essential for bacterial virulence (e.g., the pMT1, pPCP1, and pCD1 plasmids and the unstable pigmentation [*pgm*] locus, required for iron acquisition by bacteria from the host). Several are derivatives of known virulent strains (e.g., KIM, Yokohama). Fourteen *Yersinia* isolates, representing 6 non-*Y*. *pestis* species (**Table 1**), were provided by University of Florida (Drs. Tamara Revazishvili and John Glenn Morris) and included 4 *Y. pseudotuberculosis* strains and 2 strains each of *Y. aldovae*, *Y. enterocolitica*, *Y. frederiksenii*, *Y. kristensenii*, and *Y. mollaretii*.

A representative panel of “core microbiome” species of the gastrointestinal (GI) tract was selected based on several studies comparing diverse healthy individuals (64–66). The main limitation when selecting the strains was the ability of the particular isolate to be cultured, as many Bacteroidetes and/or Firmicutes have never been cultured. Therefore, several species of common culturable bacterial genera from these phyla were identified and used. *E. coli* strains were selected, if they were isolated from healthy humans and not known to be associated with diseases, to represent members of the phylum Proteobacteria. These bacteria represent “healthy gut” *E. coli*. Common probiotic bacteria, such as *Bifidobacterium* spp. found in yogurts, are also associated with the healthy microbiome (phylum Actinobacteria) and were included as part of the panel. Many of the isolates from healthy human guts are also part of the BEI Resources Human Microbiome Project initiative, and the genomes of several have been sequenced by the Broad Institute. In all, 48 isolates representing 13 genera were chosen to represent the core microbiome of the GI tract (**Table S1**).

*Challenge strain*. A fully virulent human pneumonic plague isolate, *Y. pestis* strain CO92 (biovar orientalis), was obtained from BEI Resources. *Y. pestis* CO92 was grown on tryptic soy broth (TSB) for all *in vitro* assays. Assay specific growth conditions are described below. Cultures of *Y. pestis* CO92 were grown on plates at 28°C on the indicated medium (TSB, heart infusion broth [HIB], or sheep’s blood agar [SBA]). All studies involving *Y. pestis* CO92 were performed in Biosafety level (BSL)-3, CDC-approved Tier 1 select agent laboratories at the University of Texas Medical Branch (UTMB) in the Galveston National Laboratory (GNL), Galveston, TX. Prior to the initiation of the study, the standard *Y. pestis* phenotypes for strain CO92 were confirmed by growth on various differential media, and PCR on the well characterized *Y. pestis* virulence genes, *pla* (plasminogen activating protease), *caf*1 (capsular antigen F1), and *lcrV* (low calcium response V antigen), using genomic DNA (67, 68).

### YPP-401 Bacteriophage Cocktail

YPP-401 is an aqueous, buffered, cocktail comprised of approximately equal concentrations of four different monophages (23, 28) grown on *Y. pseudotuberculosis* and was stored at 4°C protected from light until use. The YPP-401 formulation and monophage identities are proprietary to Intralytix, Inc., a company developing YPP-401 as a biothreat medical countermeasure.

### *In Vitro* Phage Susceptibility Testing

Susceptibility of the challenge strain, *Y. pestis* CO92, to the YPP-401 cocktail was assessed using the same bacterial stock used for the *in vivo* challenge studies. *Y. pestis* CO92 was grown in TSB at 28°C overnight (O.N.; ∼16 h) from a -80°C stock. The next morning, cultures were diluted 1:10 and grown for 1 h at 37°C (OD_600_ 0.3±0.15). TSB containing 0.7% agar was autoclaved and kept in a 52°C water bath until use. Phage susceptibility assays were performed using the classical spot test (29) and liquid growth assays.

For the spot tests, the bacterial culture (100 μL) was combined with 3-5 mL of 0.7% top agar and overlaid on top of 1.5% bottom agar plates. Overlay agar in plates was allowed to solidify. Serial dilutions (up to 10^-10^) of the phage cocktail were prepared. Each serial phage dilution was spotted (10 μL) individually onto the overlay plates. Plates were then incubated up to 48 h at the appropriate conditions for each bacterial species (listed in **Tables 1** and **S1**), and then examined for zones of lysis. For the liquid-based phage susceptibility assay, 200 μL of an O.N. *Y. pestis* culture was combined with 800 μL of each serial phage cocktail dilution in duplicate and incubated at 28°C or 37°C, shaking at 180 RPM, for 16-24 h. Bacterial growth was measured using turbidity (OD_600_).

### *In Vitro* Antibiotic Susceptibility Testing

The antibiotic susceptibility testing was performed using the BD BBL™ Sensi-Disc™ Antimicrobial Susceptibility Test Discs (Becton, Dickinson and Co., Franklin Lakes, NJ) per the manufacturer’s instructions. Briefly, for each isolate, a suspension of colonies was prepared by selecting several colonies from the agar plate. The suspension was diluted, as required, to obtain turbidity equivalent to 0.5 McFarland turbidity standard. A sterile cotton swab was dipped into the inoculum and then streaked across the entire agar surface of the appropriate medium plates. The plates were allowed to dry, and then the appropriate discs were aseptically applied to the surface of the agar. The plates were incubated agar side up at the appropriate conditions for 16-24h. The diameters of the zones of complete inhibition were measured to the nearest whole millimeter.

### Rat Pneumonic Plague Challenge Model

*Animals.* Brown Norway rats (*Rattus norvegicus*; 10-12 weeks old) were purchased from Charles River Laboratories (Houston, TX). Animals were acclimated for a minimum of 7 days in the Association for Assessment and Accreditation of Laboratory Animal Care International (AAALAC)-accredited, CDC-approved UTMB animal BSL-3 facility (ABSL-3) before the start of the experiment. Animals were provided food and water *ad libitum* and were maintained on a 12 h light dark cycle. Equal numbers of male (200-250 g) and female rats (100-150 g) of the same age were used in each experiment.

*Aerosol Challenge.* Rats were challenged as previously described (27) using a fully automated computer-controlled rodent whole-body aerosol exposure system (Biaera AeroMP; Biaera Technologies, Hagerstown, MD). *Y. pestis* CO92 was prepared fresh prior to each challenge as previously described (27, 69). Briefly, the strain was streaked from a -80°C stock onto an SBA plate and incubated at 28°C for 48 h. The plate was then scraped and the culture resuspended in 4 mL of HIB. The culture was diluted 1:350 in 100 ml of HIB containing 0.2% xylose in a 500 ml HEPA-filter-capped flask, and incubated for 24 h at 30°C, shaking at 100 RPM. The culture was then washed twice in HIB and resuspended in 1/10^th^ the original culture volume (i.e., 10 mL) of HIB. A further 1:10 dilution was performed, and Antifoam A Concentrate (Sigma-Aldrich, Saint Louis, MO) was added to a final concentration of 0.2%. The aerosol challenge preparation was then titrated on SBA plates after 10-fold serial dilutions to confirm bacterial concentration.

Separate aliquots were prepared for each aerosol exposure run since only 8 rats could be accommodated in a single run. Aliquots of HIB (20 mL) supplemented with 4% glycerol and 0.2% Antifoam A were prepared for each run to quantify bacterial numbers in samples collected by an SKC BioSampler® (SKC, Inc., Eighty Four, PA). The following aerosol parameters were used: an air inflow rate of 30L/min, with 14L/min to the nebulizer and 16L/min to dilute the air; and an air outflow rate of 30L/min, with 12.5L/min to the bio-sampler and 17.5L/min to the exhaust. The humidity was maintained at >60% and the aerosol exposure time of *Y. pestis* to animals was 15 min using a 6-jet Collison nebulizer (CH Technologies [USA], Inc., Westwood, NJ). An Aerodynamic Particle Sizer (TSI) was used to validate the aerosol delivery equipment, specifically the aerosolization efficiency (spray factor; SF), aerosol concentration derived from the samples (CSAMP), and the concentration of bacteria in the suspension pipetted into the nebulizer (CNEB). Due to practical limitations on number of animals that could be exposed to bacteria at one time, the challenge was divided into staggered consecutive runs. After bacterial exposure, rats were randomly distributed among treatment groups to account for any potential variability in presented dose (Dp) between each run. The input bacterial concentration was 1 x 10^9^ CFU/mL with a Dp to rats ranging from 9.56 x 10^6^ to 1.24 x 10^7^ CFUs for all runs.

*Dosing Regimen.* As indicated earlier, phage treatment was initiated at either 18 or 42 hpi. The levofloxacin antibiotic positive control (Akorn, Inc., Lake Forest, IL; NDC: 17478-107-20) was administered once daily at 50 mg/kg via the oral route for 10 days. The vehicle (i.e., placebo; 100 μL 1X phosphate buffered saline [PBS]) control was administered intraperitoneally (i.p.) starting at either 18 or 42 hpi, and as indicated. Oral administration of the phage preparation was perform as described (31) using disposable animal feeding needles (20G × 1.50 in.; Thermo Fisher Scientific, location!!). The volumes of each dose of YPP-401 for various routes were: 500 μL oral; 500 μL i.p.; 500 μL i.m. at 250 μL per hind leg; and 100 μL i.n. at 50 μL per naris under anesthesia via inhalation of isofluorane. When treatment was initiated at 18 hpi, vehicle and YPP-401 were administered by the indicated route four times over 2 days (approximately every 6-12 h) at 18, 24, 30 and 42 hpi. When treatment was initiated at 42 hpi, vehicle and YPP-401 were intended to be administered by the indicated route four times over 2 days (approximately every 6-12 h) at 42, 48, 66 and 72 hpi. Number of doses actually administered when treatment was initiated at 42 hpi are indicated in **Table S4**.

*Efficacy.* Infected and treated rats were observed for morbidity and mortality twice per day for up to 14-15 days with 3 checks/day on days 3-6 post-infection (dpi) or when animals exhibited increased clinical scores. Clinical outcome measures included rate of survival at 14 dpi, weight loss (measured every other day during 0-7dpi and 2X per week thereafter), and clinical scores. Clinical scores were quantitative assessments using an independent variable score for appearance, activity level, respiration, and facial expression. The rodent clinical scoring rubric was approved by the UTMB Institutional Animal Care and Use Committee (IACUC).

### Quantification of Bacterial Load

Aerosol samples were collected in a SKC BioSampler and were ten-fold serially diluted and spotted on SBA plates to quantify bacterial numbers. To quantify bacteremia, the blood was collected at 42 hpi (and prior to initiation of treatment for animals dosed at 42 hpi). Organs/tissues (i.e., lung, liver, heart, spleen) were collected from moribund rats at time of euthanasia or from surviving rats at 14-15 dpi (i.e., end of the study). Tissues were homogenized by placing lung, spleen, or heart into 15 mL Closed Tissue Grinder Systems (Fisherbrand, Pittsburgh, PA) with 2 mL of Dulbecco’s PBS (DPBS). Liver was homogenized using 50 mL Closed Tissue Grinder System with 4 mL DBPS. Blood and tissue homogenates were 10-fold serially diluted in DPBS and then plated onto SBA plates to quantify bacterial load.

### Statistical methods

Statistical analysis was performed using GraphPad Prism version 10.0.0 for Windows (GraphPad Software, Boston, MA). To compare the *Y. pestis* burden in blood at 42hpi between treatment groups, a one-way analysis of variance (ANOVA) was conducted for each cohort, followed by Tukey’s post-hoc analysis for multiple comparisons. To compare the terminal bacterial burden in each organ between treatment groups, a two-way analysis of variance (ANOVA) was conducted for each cohort, followed by Tukey’s post-hoc analysis for multiple comparisons for each organ.

## ACKNOWLEDGEMENTS

This project has been funded in whole or in part with Federal funds from the Division of Microbiology and Infectious Diseases, National Institute of Allergy and Infectious Diseases, National Institutes of Health, Department of Health and Human Services, under Contract No. HHSN272201700040I/75N93022F00003.

We thank Dr. Joseph Campbell (DMID/NIAID) for guidance and Drs. Tamara Revazishvili and John Glenn Morris (University of Florida) for their contributions to this work. Thanks are also extended to Dr. David W. Beasley, Jennifer Young, and Sudjai W. Montgomery, UTMB, for providing administrative oversight on the project.

P.B.K was supported in part by the Texas Medical Center Training Program in Antimicrobial Resistance (TPAMR) T32 Fellowship Program AI 141349.

The following authors contributed to each of the following areas: Conceptualization and study design: A.K.C., A.S., J.A.S., J.S., J.W.; Formal analysis: J.A.S., J.W., P.B.K. J.S.; Funding acquisition: A.K.C., A.S., J.A.S.; Investigation: L.H., J.S., P.B.K., E.K.H., B.H.M., W.S.L., J.E.P.; Methodology: A.K.C., J.S,, P.B.K., E.K.H., B.H.N., W.S.L., J.E.P.; Administration and supervision: A.K.C., A.S., J.A.S., J.S.; Resources: J.W., A.K.C., L.H.; Writing: P.B.K., J.A.S.; Editing: A.K. C., J.S., P.B.K., A.S., J.W. All authors have reviewed the manuscript.

## SUPPLEMENTARY TABLES

**Table S1.** Representative Non-Yersinia Host Microbiome Isolates for Specificity Testing.

| Phylum | Species | Source / Strain |
| --- | --- | --- |
| Actinobacteria | *Bifidobacterium angulatum* | BEI HM-1189 |
|  | *Bifidobacterium breve* | BEI HM-856 |
|  | *Bifidobacterium longum* | ATCC 55813 |
|  | *Bifidobacterium longum* | ATCC 55814 |
|  | *Bifidobacterium longum* | ATCC BAA-999 |
|  | *Bifidobacterium* sp. | BEI HM-30 |
|  | *Bifidobacterium/Lactobacillus* sp. | ATCC 11146 |
|  | *Streptomyces* sp. | BEI HM-859 |
|  | *Streptomyces* sp. | BEI HM-789 |
| Bacteroidetes | *Bacteroides caccae* | BEI HM-728 |
|  | *Bacteroides dorei* | BEI HM-717 |
|  | *Bacteroides fragilis* | ATCC 700786 |
|  | *Bacteroides fragilis* | BEI HM-710 |
|  | *Bacteroides fragilis* | BEI HM-20 |
|  | *Bacteroides fragilis* | BEI HM-714 |
|  | *Bacteroides salyersiae* | BEI HM-725 |
|  | *Bacteroides* sp. | BEI HM-18 |
|  | *Bacteroides* sp. | BEI HM-23 |
|  | *Bacteroides* sp. | BEI HM-22 |
|  | *Bacteroides* sp. | BEI HM-19 |
|  | *Parabacteroides merdae* | ATCC 43184 |
|  | *Prevotella nigrescens* | BEI HM-1053 |
|  | *Prevotella oralis* | BEI HM-1054 |
| Firmicutes | *Clostridium innocuum* | BEI HM-173 |
|  | *Enterococcus faecalis* | ATCC BAA-2820 |
|  | *Enterococcus faecalis* | ATCC 51299 |
|  | *Enterococcus faecalis* | BEI HM-432 |
|  | *Enterococcus faecalis* | BEI HM-202 |
|  | *Enterococcus faecalis* | BEI NR-32002 |
|  | *Enterococcus faecium* | BEI HM-975 |
|  | *Enterococcus faecium* | BEI HM-952 |
|  | *Enterococcus faecium* | BEI HM-959 |
|  | *Enterococcus faecium* | BEI HM-204 |
|  | *Lactobacillus brevis* | ATCC 14869 |
|  | *Ruminococcus gnavus* | BEI HM-1056 |
|  | *Ruminococcus lactaris* | BEI HM-1057 |
|  | *Ruminococcus* sp. | BEI HM-79 |
|  | *Ruminococcus torques* | ATCC 27756 |
|  | *Ruminococcus torques* | ATCC 35915 |
|  | *Streptococcus parasanguinis* | BEI HM-1060 |
|  | *Streptococcus sanguinis* | BEI HM-1061 |
| Fusobacteria | *Fusobacterium ulcerans* | BEI HM-1194 |
| Proteobacteria | *Escherichia coli* | BEI HM-366 |
|  | *Escherichia coli* | BEI HM-365 |
|  | *Escherichia coli* | BEI HM-347 |
|  | *Escherichia coli* | BEI NR-17674 |
|  | *Escherichia coli* | BEI NR-17676 |
|  | *Pseudomonas* sp. | BEI HM-214 |

**Table S2.**
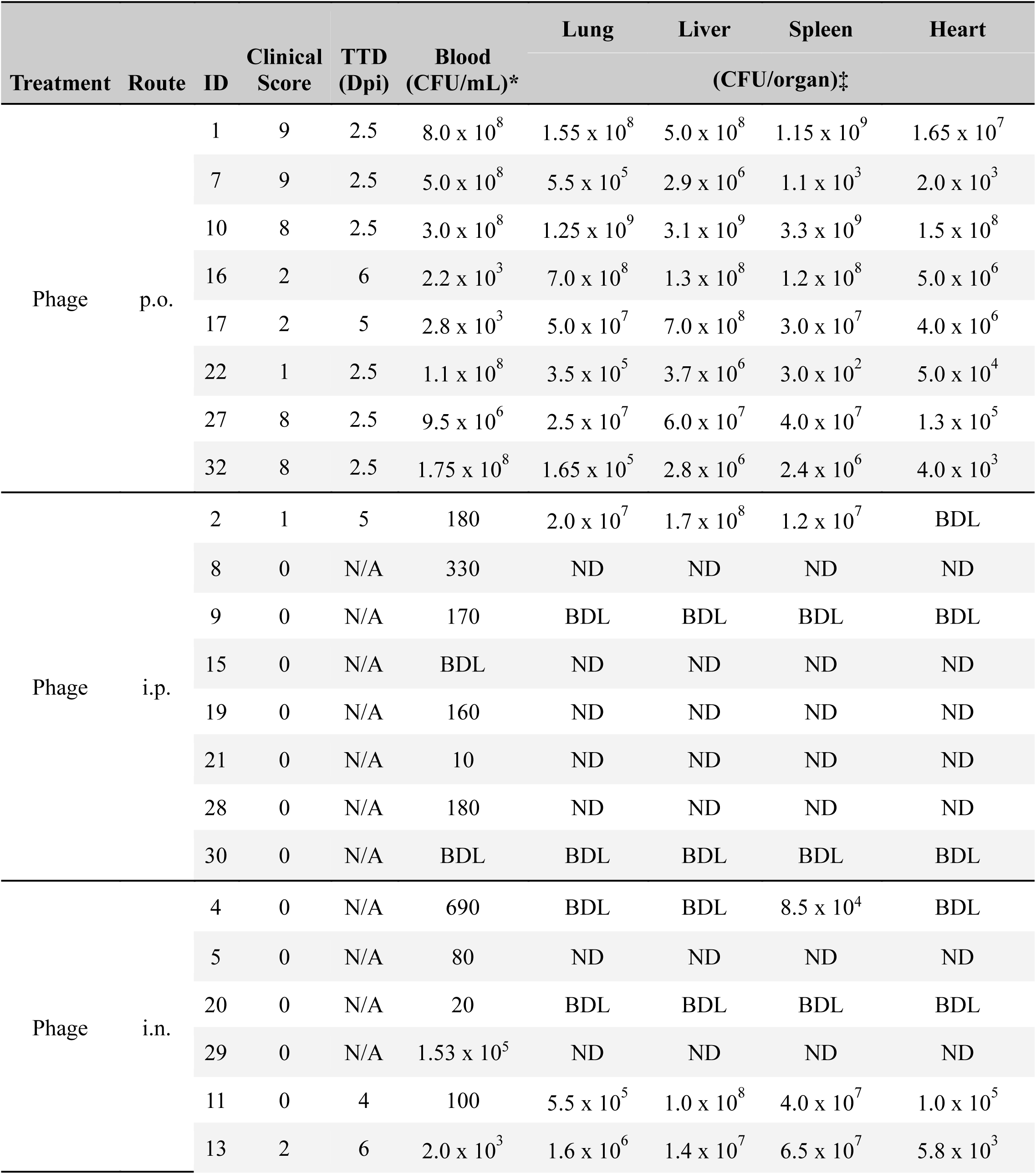
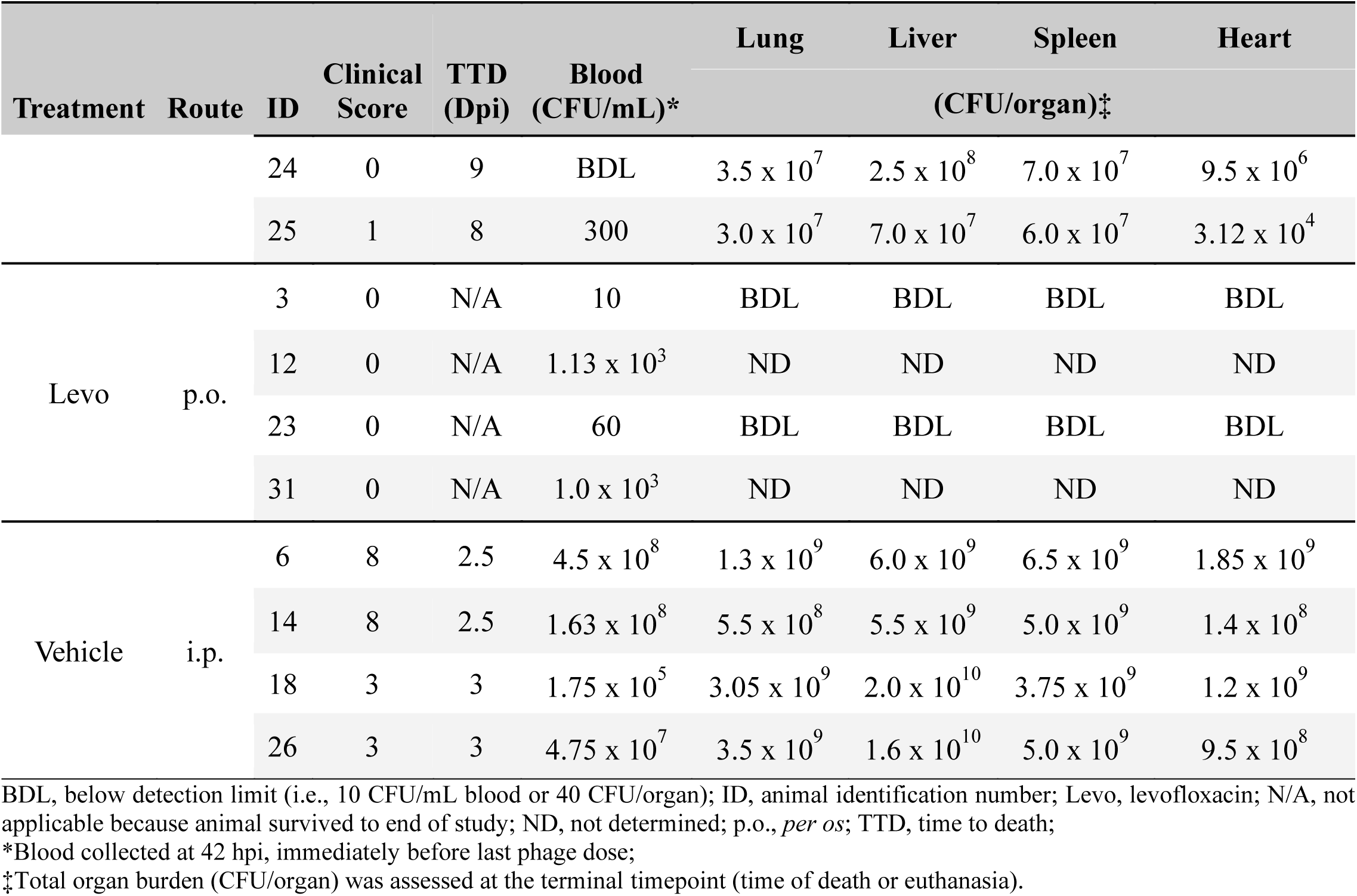
Clinical Measures and Bacterial Burden in Cohort 1 Rats Challenged at 18 hpi.

**Table S3.** Clinical Measures and Bacterial Burden in Cohort 2 Rats Challenged at 18 hpi.

| Treatment | Route | ID | Clinical Score¶ | TTD (Dpi) | Blood (CFU/mL)* | Lung | Liver | Spleen | Heart |
| --- | --- | --- | --- | --- | --- | --- | --- | --- | --- |
|  |  |  |  |  |  | (CFU/organ)‡ |  |  |  |
| Phage | i.n. | 1 | 0 | N/A | BDL | BDL | BDL | BDL | BDL |
|  |  | 6 | 0 | N/A | BDL | BDL | BDL | BDL | BDL |
|  |  | 11 | 0 | N/A | BDL | 2.1 x 10<sup>6</sup> | BDL | BDL | 6.5 x 10<sup>4</sup> |
|  |  | 16 | 0 | N/A | 5.0 x 10<sup>3</sup> | BDL | BDL | BDL | BDL |
|  |  | 22 | 0 | N/A | BDL | BDL | BDL | BDL | BDL |
|  |  | 27 | 0 | N/A | BDL | BDL | BDL | BDL | BDL |
|  |  | 32 | 0 | N/A | 5.0 x 10<sup>2</sup> | BDL | BDL | BDL | BDL |
|  |  | 17 | 0 | 6 | BDL | 4.5 x 10<sup>2</sup> | 4.9 x 10<sup>7</sup> | 3.1 x 10<sup>8</sup> | 4.0 x 10<sup>3</sup> |
| Levo | p.o. | 7 | 0 | N/A | BDL | BDL | BDL | BDL | BDL |
|  |  | 18 | 0 | N/A | BDL | BDL | BDL | BDL | BDL |
| Vehicle | i.p. | 2 | 9 | 2.5 | 1.65 x 10<sup>7</sup> | 5.0 x 10<sup>8</sup> | 1.2 x 10<sup>9</sup> | 1.25 x 10<sup>9</sup> | 1.95 x 10<sup>8</sup> |
|  |  | 21 | 7 | 3 | 3.0 x 10<sup>6</sup> | 2.0 x 10<sup>6</sup> | 8.0 x 10<sup>3</sup> | 2.0 x 10<sup>4</sup> | 4.5 x 10<sup>3</sup> |

BDL, below detection limit (i.e., 10 CFU/mL blood or 40 CFU/organ); ID, animal identification number; Levo, levofloxacin; N/A, not applicable because animal survived to end of study; ND, not determined; p.o., *per os*; TTD, time to death;
¶ Clinical score at time of death or euthanasia;
\*Blood collected at 42 hpi, immediately before last phage dose;
‡ Total organ burden (CFU/organ) was assessed at the terminal timepoint (time of death or euthanasia).

**Table S4.** Clinical Measures and Bacterial Burden in Cohort 3 Rats Challenged at 42 hpi.

| Treatment | Route | ID | #<br>Doses† | Clinical<br>Score¶ | TTD<br>(Dpi) | Blood<br>(CFU/mL)* | Lung | Liver | Spleen | Heart |
| --- | --- | --- | --- | --- | --- | --- | --- | --- | --- | --- |
|  |  |  |  |  |  |  | (CFU/organ)‡ |  |  |  |
| Phage | i.m. | 3 | 2 | 7 | 3 | $7.5 \times 10^5$ | $1.6 \times 10^8$ | $1.9 \times 10^9$ | $9.5 \times 10^8$ | $1.05 \times 10^8$ |
| | | 8 | 1 | 3 | 2.5 | $5.0 \times 10^7$ | $3.0 \times 10^6$ | $4.0 \times 10^6$ | $4.0 \times 10^6$ | $1.5 \times 10^6$ |
| | | 10 | 2 | 7 | 3 | $2.5 \times 10^4$ | $5.5 \times 10^6$ | $1.7 \times 10^7$ | $1.5 \times 10^6$ | $1.0 \times 10^6$ |
| | | 13 | 1 | 2 | 2.5 | $6.0 \times 10^7$ | $2.0 \times 10^6$ | $4.0 \times 10^6$ | $4.5 \times 10^6$ | $1.5 \times 10^6$ |
| | | 19 | 2 | 7 | 3 | $1.75 \times 10^5$ | $3.0 \times 10^8$ | $5.0 \times 10^9$ | $7.5 \times 10^8$ | $1.1 \times 10^8$ |
| | | 24 | 2 | 7 | 3 | $4.0 \times 10^5$ | $8.5 \times 10^6$ | $1.7 \times 10^7$ | $3.5 \times 10^6$ | $5.5 \times 10^6$ |
| | | 25 | 2 | 7 | 3 | $1.0 \times 10^5$ | $2.5 \times 10^6$ | $2.2 \times 10^8$ | $4.5 \times 10^7$ | $2.0 \times 10^6$ |
| | | 30 | 2 | 7 | 3 | $4.0 \times 10^7$ | $1.5 \times 10^6$ | $1.1 \times 10^6$ | $2.5 \times 10^5$ | $7.0 \times 10^4$ |
| Phage | i.p. | 4 | 2 | 7 | 3 | $7.0 \times 10^5$ | $1.9 \times 10^8$ | $1.4 \times 10^9$ | $6.0 \times 10^8$ | $1.8 \times 10^8$ |
| | | 5 | 1 | 1 | 2.5 | $1.5 \times 10^7$ | $4.5 \times 10^6$ | $7.0 \times 10^6$ | $1.3 \times 10^7$ | $1.0 \times 10^6$ |
| | | 9 | 1 | 0 | 2.5 | $5.0 \times 10^7$ | $3.5 \times 10^6$ | $8.0 \times 10^6$ | $7.5 \times 10^6$ | $1.5 \times 10^6$ |
| | | 14 | 1 | 3 | 2.5 | $4.5 \times 10^7$ | $3.5 \times 10^5$ | $4.8 \times 10^4$ | $4.0 \times 10^3$ | $3.0 \times 10^3$ |
| | | 20 | 1 | 9 | 2.5 | $1.0 \times 10^5$ | $7.5 \times 10^4$ | $1.0 \times 10^3$ | $4.0 \times 10^3$ | $4.5 \times 10^2$ |
| | | 23 | 2 | 7 | 3 | $5.5 \times 10^5$ | $6.0 \times 10^7$ | $1.3 \times 10^9$ | $6.0 \times 10^8$ | $7.0 \times 10^7$ |
| | | 28 | 2 | 7 | 3 | $1.4 \times 10^6$ | $6.0 \times 10^7$ | $1.6 \times 10^9$ | $1.25 \times 10^7$ | $4.0 \times 10^7$ |
| | | 29 | 2 | 7 | 3 | $5.0 \times 10^3$ | $7.5 \times 10^5$ | $1.7 \times 10^5$ | $5.0 \times 10^3$ | $6.0 \times 10^2$ |
| Vehicle | i.p. | 12 | 1 | 9 | 2.5 | $1.3 \times 10^7$ | $4.0 \times 10^9$ | $7.0 \times 10^9$ | $7.0 \times 10^9$ | $5.0 \times 10^8$ |
| | | 31 | 2 | 7 | 3 | $7.5 \times 10^6$ | $7.5 \times 10^8$ | $6.0 \times 10^9$ | $3.5 \times 10^9$ | $3.0 \times 10^8$ |
| Levo | p.o. | 15 | 10 | 0 | N/A | $6.5 \times 10^4$ | BDL | BDL | BDL | BDL |
| | | 26 | 1 | 7 | 3 | $2.5 \times 10^7$ | $3.0 \times 10^3$ | $1.7 \times 10^4$ | $4.0 \times 10^4$ | $3.5 \times 10^2$ |

BDL, below detection limit (i.e., 10 CFU/mL blood or 40 CFU/organ); ID, animal identification number; Levo, levofloxacin; N/A, not applicable because animal survived to end of study; ND, not determined; p.o., *per os*; TTD, time to death;
† Total number of treatment doses each animal received prior to euthanasia or succumbing to death;
¶ Clinical score at time of death or euthanasia;

| Treatment | Route | ID | #<br>Doses† | Clinical<br>Score¶ | TTD<br>(Dpi) | Blood<br>(CFU/mL)* | Lung | Liver | Spleen | Heart |
| --- | --- | --- | --- | --- | --- | --- | --- | --- | --- | --- |
|  |  |  |  |  |  |  | (CFU/organ)‡ |  |  |  |

\* Blood collected at 42 hpi, immediately before first phage dose;
‡ Total organ burden (CFU/organ) was assessed at the terminal timepoint (time of death or euthanasia).

---

**Figure S1.**
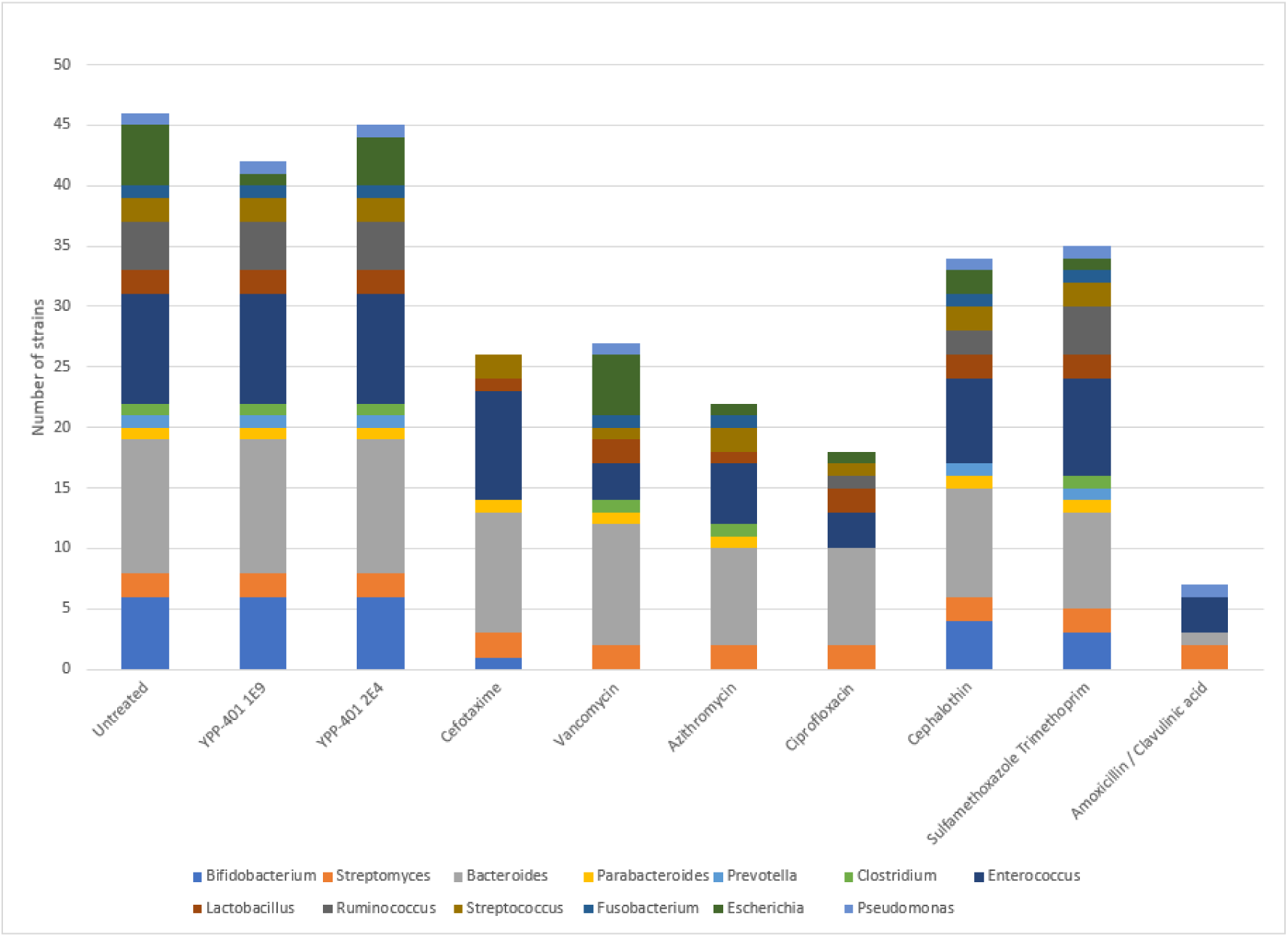
YPP-401 is Highly Specific for *Yersinia* spp. Compared to Antibiotics. Susceptibility testing was performed on the indicated bacterial species (**Table S2**). Susceptibility to YPP-401 at two concentrations was assessed using spot and/or liquid growth assays. Susceptibility to the indicated antibiotic was performed using the BD BBL™ Sensi-Disc™ Antimicrobial Susceptibility Test Discs (Becton, Dickinson and Co., Franklin Lakes, NJ).

